# Enhancing In Vitro SMN Protein Expression and Cell Viability through Xeno-Nucleic Acid-Based ASOs in Spinal Muscular Atrophy

**DOI:** 10.1101/2023.08.21.552929

**Authors:** Ozum Kilic, Hale Ahsen Babar, Cemre Can Inci, Sibel Pinar Odabas, Gamze Yelgen, Sevgi Oltan, Sila Kulac, Cihan Tastan

**Affiliations:** Molecular Biology, Institute of Science and Technology, Üsküdar University, Istanbul, TURKEY; Transgenic Cell Technologies and Epigenetic Application and Research Center (TRGENMER), Üsküdar University, Istanbul, TURKEY; HiDNA Biotechnology Inc., Istanbul, TURKEY

**Keywords:** Spinal muscular atrophy, Survival Motor Neuron, Xeno Nucleic Acid, Antisense Oligonucleotide, ASO

## Abstract

Spinal Muscular Atrophy (SMA) stands as a devastating ailment arising from the dearth of functional SMN (Survival Motor Neuron) protein due to genetic anomalies within the SMN1 gene. This condition is marked by the consequential attrition of motor neurons, precipitating a progressive decline in muscular strength and culminating in the disruption of neuromuscular junctions. Existing therapeutic approaches encompassing Zolgensma, Nursinersen, and Evrysdi employ innovative genetic therapeutic strategies involving transgene delivery, Antisense Oligonucleotide (ASO) technology, and modulation of pre-mRNA processing to enhance functional SMN protein expression. However, the ASO therapeutics remain suboptimal in establishing a sustained panacea for SMA, as they inadequately maintain consistent levels of functional SMN protein expression. In this study, we present a discerning inquiry into focusing on XNA-DNA-ASO products that exhibit enhanced safety and stability compared to conventional DNA/RNA-ASO sequences. Through precise targeting of the ISSN-1 region within SMN2 gene’s intron 7, our approach seeks to amplify SMN protein expression. Employing Xeno Nucleic Acid (XNA) bases, known for their augmented hydrophobicity and stability, our strategy surmounts previous limitations associated with chemical modifications, showcasing heightened endonuclease resistance. Comparative analyses with conventional DNA/RNA-ASO products substantiate the superiority of XNA-DNA-ASO sequences, underscoring elevated SMN protein expression and reduced toxicity. In a comprehensive evaluation, our gene therapy paradigm is scrutinized within a type 1 SMA fibroblast cell line. Utilizing diverse analytical methodologies, encompassing Annexin V-PI analysis for cytotoxicity, MTT assays for mitochondrial activity, and flow cytometry for SMN protein expression profile, we gauge therapeutic impact and potential toxicity. In conclusion, our investigation not only spotlights the promise of XNA-DNA-ASO sequences but also holds implications for refining SMA treatment strategies, converging on minimized dosages, lowered toxicity, and heightened therapeutic efficacy, thus shaping the landscape of gene therapy for SMA.

## Introduction

Spinal Muscular Atrophy (SMA) stands as a profound neuromuscular affliction marked by the progressive demise of motor neurons and muscular atrophy, emanating from underlying genetic anomalies. The crux of SMA pathology is grounded in the inability to generate fully functional SMN (Survival Motor Neuron) protein due to diverse deletions and mutations in both alleles of the Survival Motor Neuron 1 (SMN1) gene (Singh et al., 2017). Reverberating across various cellular processes including snRNP biogenesis, transcription, translation, stress granule formation, and macromolecular trafficking, SMN protein emerges as an indispensable housekeeping entity (Singh et al., 2017). Its absence culminates in an irreversible attrition of anterior horn cells in lower motor neurons and brainstem nuclei within the spinal cord, engendering augmented muscle debilitation (Prior et al., 2009). SMA exhibits a substantial heterogeneity of SMN1 gene mutations, with exon 7 deficiency in the SMN protein affecting approximately 95% of SMA patients (Wirth, 2000). This pivotal segment spanning 16 amino acids encoded by exon 7 bears essential implications for SMN protein stability and oligomerization, underpinning its functional conformation (Lorson et al., 1999). The paradigmatic influence of the SMN2 gene, yielding only ∼10% full-length (FL) SMN protein in contrast to the 100% produced by the SMN1 gene in healthy individuals, dovetails into the phenotypic severity of SMA and the correlation between SMN2 copy numbers and disease manifestation (Bowerman et al., 2017).

The therapeutic landscape for SMA encompasses gene editing methods, epitomized by “Zolgensma (Onasemnogene abeparvovec),” an adenoviral vector-mediated delivery of the SMN1 gene primarily for SMA type 1 infants (Hoy, 2019; Mendell et al., 2017). Yet, Onasemnogene abeparvovec’s transient impact on SMN1 gene expression and concerns regarding sustained efficacy curtail its promise (Lin et al., 2020). “Spinraza (Nusinersen),” an ASO-based strategy, modulates exon 7 inclusion within SMN2 gene, while “Evrysdi (Risdiplam)” intervenes in pre-mRNA processing, both avenues bolstering functional SMN protein production (Wurster and Ludolph, 2018; Messina and Sframeli, 2020). Within this context, Nusinersen’s mode of action centers on mitigating secondary structures like stem loops within the ISS-N1 region of SMN2, thereby relieving silencer elements and enhancing exon 7 incorporation (Singh et al., 2013; Singh et al., 2015).

Antisense oligonucleotide (ASO) technology, as a steadfast alternative to strategies imposing permanent genome modifications, emerges as a cost-effective and steadfast alternative (Hagedorn et al., 2017). Nusinersen, the inaugural therapeutic for SMA, emerges as a potent contender in thwarting the formation of a secondary stem loop within the ISS-N1 region (positions 10 and 24 in intron 7) of the SMN2 gene, by meticulously targeting this domain with its ASO sequences (Singh et al., 2013; Singh et al., 2015). This secondary conformation, known as the stem loop, crystallizes through the interaction between the cytosine (C) at position 10 of the ISS-N1 region and the guanine (G) located at position 290. These internal stem loop structures (ITSL) offer anchorage to silencer elements—hnRNPA1/A2B1 proteins—thereby impeding the engagement of factors like T1A1, which stimulate U1 snRNP proteins, with uridine-rich regions (URC-1/2). Consequently, U1 snRNP’s ability to effectively recognize exon 7, crucial for SMN protein expression, is hindered, culminating in a meager expression level of full-length SMN protein (∼10%) (Singh et al., 2007; Singh et al., 2015). Intriguingly, compensatory mutations strategically introduced within intron 7 or exon 7, aimed at attenuating or obstructing the formation of these secondary structures, wield profound influence on augmenting SMN2 exon 7 incorporation, hence elevating functional SMN protein expression.

Amidst the architectural scaffolds of natural DNA and RNA, the landscape has witnessed the emergence of hydrophobic Xeno-Nucleic Acid (XNA) polymers, showcasing diverse chemical backbones (Chaput et al., 2019). These XNA molecules embody variations in sugar moieties, yet retain nucleobases akin to their DNA and RNA counterparts. Notably, XNA sequences boast an elevated affinity for binding targeted RNA sequences and evince enhanced resistance against native nuclease enzymes (Schmidt M., 2010). Harnessing these attributes, XNA-ASO ensembles usher in RNA-based therapeutic strategies, offering a nuanced avenue characterized by diminished toxicity (Hagedorn et al., 2017). Additionally, the potential to modulate splicing mechanisms surfaces as XNA sequences, when incorporated at the extremities of DNA sequences, create hybrid XNA-DNA mixmer structures with implications for splicing mechanics. Moreover, XNA probes demonstrate their capability to forge robust complexes through high-specificity binding to intended DNA or RNA sequences (Hagedorn et al., 2017). Collectively, the emergence of XNA sequences promulgates a paradigm shift, presenting next-generation biopolymers endowed with distinctive chemical attributes that hold the potential to supplant traditional DNA and RNA, thereby ushering diverse research and therapeutic prospects. Within our investigatory focus on intron 7, the crux of the matter revolves around the pre-mRNA splicing machinery, delineating a critical bottleneck wherein the SMN2 gene, a cornerstone in fulfilling the functional SMN protein requisites for SMA patients, languishes in its ability to engender substantive SMN protein levels. To address this challenge, our study pivots on a pivotal question: how can a method be devised that caters universally to all SMA patients? In delineating the path of genetic therapy, a myopic approach concentrating solely on a single SMN1 gene mutation limits the scope of patients amenable to treatment. Thus, the salient avenue lies in strategies attuned to the SMN2 gene, which shares an astonishing 99% homology with SMN1. By championing the potential of this avenue, our research serves as a beacon illuminating an extended spectrum of patients enfranchised by the prospective treatments that our inquiry may foster. This calls for the exploration of intron 7-targeted next-generation paradigms poised to augment SMN2 gene engagement, thereby heralding the promise of sustainable treatments for a panoply of genetic maladies, including SMA.

Our present study pivots on XNA-DNA-ASO products, a distinct subclass tailored to target the ISS-N1 region nested within intron 7 of the SMN2 gene. This novel approach, distinct from prevalent DNA-ASO counterparts, is imbued with potential to redefine the landscape of new-generation genetic therapies. Implemented within the milieu of SMA Type I fibroblast cell line, our study unveils novel dimensions of next-generation genetic interventions. Deliberately designed XNA/DNA-ASO mixmers leverage strengthened H-bonds, whether double (A-T) or triple (G-C), to transcend the limitations of first-generation DNA-ASOs, particularly in ISS-N1 region-specific bases. Central to our pursuit is the assessment of these designed mixmers’ relative merits in terms of toxicity and the quantum of SMN protein expression. Herein, we present two specific DNA/XNA-ASO sequences tailored for the ISS-N1 target site, alongside a DNA-ASO sequence featuring 2′-O-methoxyethyl (MOE) modification and phosphorothioate structure, which serves as a positive control benchmarked against prevailing ASO treatment strategies. Guided by these considerations, we identify the optimum XNA-DNA-ASO sequence for ISS-N1 that maximizes exon 7 involvement with remarkable efficacy, thereby selecting a candidate for further validation through trials on SMA Type I fibroblast cell line. By monitoring resultant changes in SMN protein expression, our research not only charts new avenues for probing novel generation XNA-DNA-ASO gene therapy strategies but also advances potential interventions that validate their capacity to elevate functional SMN protein expression levels in an in vitro milieu.

This study explores novel frontiers by focusing on XNA-DNA mixmer polymers, alternate to natural deoxyribose and ribose sugars, with potential for low-toxicity RNA-based therapies (Chaput et al., 2019; Hagedorn et al., 2017). These XNA molecules confer increased stability and binding affinity, thus holding promise for a more nuanced treatment strategy. Given the exigencies of addressing a wide array of SMA patients, our research centers on the SMN2 gene, aiming to devise a universally applicable therapeutic approach. This holistic endeavor challenges the convention of focusing solely on SMN1 gene mutation. We investigate XNA-DNA-ASO sequences within the ISS-N1 region, their design tailored to maximize double H bonding and endonuclease resistance. Through meticulous assays, we ascertain their potential advantages in terms of toxicity and SMN protein expression, rendering them potential agents for enhancing functional SMN protein levels (Hagedorn et al., 2017). This endeavor ushers in a prospective era of next-generation genetic therapy strategies, replete with stability and reduced toxicity. These findings are anticipated to pave the way for in vitro gene therapy approaches demonstrating increased functional SMN protein expression, potentially transforming the therapeutic landscape of SMA.

## Material-Method

### XNA-DNA ASO mixmer Design and Delivery

Two different ASO varieties with different sequence designs were compared to determine the optimum XNA-DNA-ASO sequence. The targeted region of the XNA-DNA-ASO mixmer sequences is the ISS-N1 motif sequence within the *SMN2* intron 7 region (10-24). These XNA-DNA mixmers are designed to strengthen hydrogen bonds by grouping them separately according to the hydrogen bonds between the A-T or G-C bases they contain. As ASO sequences, phosphorothioated XNA-DNA mixmers containing DNA nucleobases (G *, A *, T *, C*) were designed for this research **(Table 1**). Using the service procurement method, ASO sequences designed to form a compact structure with the target mRNA sequence were obtained from Qiagen Biotechnology (Hilden, Germany). The designed XNA-ASO arrays were produced at 10 nmol concentration using high-performance liquid chromatography (HPLC) purification technique. The produced XNA-DNA-ASO sequences were transferred to SMA type 1 fibroblast cell line with PEI transfection reagent (Bremer, J. et al, 2022). RNA-ASO sequence with phosphorothioate modifications of 18-mer 2′-O-(2-methoxyethyl) (MOE) was designed as a confirmed Nusinersen analogue (Intron 7 position;10-27) 5′ -TCACTTTCATAATGCTGG-3′. RNA-ASO products synthesized by adhering to this modification and sequence were produced by Qiagen Biotechnology. The efficiency of the two engineered ASO sequences was controlled against the positive control RNA-ASO sequence (**Table 1**).

**Table 1:** XNA-DNA-ASO mixmer sequences targeted at SMN2 gene intron 7 region

| XNA-DNA-ASO | XNA-DNA mixmer sequence |
| --- | --- |
|  | Phosphorothioated XNA* bases/ Phosphorothioated DNA* bases |
| GENLiNA | T*+C*A*+C*T*T*T+C*A*T*A*A*T*+G*+C*T*+G*+G |
| MERLiNA | +T*C*+A*C*+T*+T*+T*C*+A*+T*+A*+A*+T*G*C*+T*G*G |
| Positive Control | mU*mC*A*mC*mU*mU*mC**A*mU*A*A*mU*G*mC*mU*G*G |

### Cell Culture

SMA Type 1 fibroblast cells (Coriell Institute, GM00232) were cultured with EMEM medium with 10% fetal bovine serum (FBS), 200 U/ml penicillin/streptomycin antibiotic, and 2 mM L-glutamine. Cells were incubated at 37°C, 5% CO_2_.

### Flow Cytometry

SMA fibroblast cells were incubated with anti-SMN-AlexaFlour 647 (Santa Cruz Biotechnology, Anti-SMN Antibody (2B1)) monoclonal antibody to determine SMN expression levels. Fixation/permeabilization procedures were performed using the BDCytofix/Cytoperm Fixation/Permeabilization Kit (Cat. No. 554714). After 100.000 cells were collected, 200 μl of Fixation/Permeabilization solution (BD) was added and incubated at 4°C for 20 minutes. After washing with BD Perm/Wash™ Buffer (BD), cells were stained with anti-SMN-AlexaFlour 647 at a dilution ratio of 1:50 and incubated at 4°C for 30 minutes. Cells were washed with BD Perm/Wash™ Buffer (BD) and analyzed by flow cytometry (Beckman Coulter, CytoFlex) to assess the percentage and the Mean Fluorescence Intensity (MFI) level of SMN expression.

### Cell Survival Analysis

Survival analysis was performed to control the cell viability of cells. Trypan blue (Biological Industries, #03-102-1B) was applied to identify and count surviving cells. Cell counting and viability analysis were performed with the BIO-RAD TC20 Automated Cell Counter.

### Toxicity Analysis

As a result of the application of the work packages to target cell lines, toxicity (apoptosis) was determined with the help of Flow Cytometry. Toxicity determination was performed according to the vendor’s instructions using the Annexin V-FITC kit (Sigma Aldrich) and Propidium iodide (Thermofisher, # P1304MP). 100 μL of cell suspension was prepared in each tube, diluting 1 × 10^5^ cells in 1X Annexin binding buffer. 5 μL of Annexin V-FITC and 2.5 μL of PI working solution (100 μg/mL) was then added to each tube. After 15 minutes of incubation at room temperature, then it was mixed with 400 μL binding buffer. Finally, flow cytometry analysis was performed by measuring the fluorescence of the stained cells at 530 nm. In addition, as a result of the application of these work packages to target cell line, their cytotoxicity, ELISA (FLUOstar, OMEGA) and MTT (3-(4,5-dimethylthiazol-2-yl)-2,5-diphenyltetrazolium bromide) tests were performed.

### Statistical Approach

Data recorded by flow cytometry were analyzed using CytExpert (Beckman Coulter). Statistical analyses were performed using GraphPad Prism 8.0.1 software (GraphPad Software, Inc., San Diego, CA, USA) with a two-tailed t-test using independent mean values. Error bars represent Standard Error of Means (SEM). For all experiments, significance was defined as p<0.05 and NS= non-Significant.

## Results

The efficacy of classical ASO sequences targeted at SMN’ to enhance exon 7 inclusion has been well-documented in numerous studies (Singh et al., 2013; Singh et al., 2015). The splicing process plays a pivotal role in modulating the functional protein production rates of the SMN2 gene as compared to its counterpart, SMN1. By strategically targeting the ISS-N1 region within intron 7, there arises an opportunity to manipulate splicing via XNA-DNA-ASO sequences, with the aim of augmenting the expression of functional SMN protein. The envisaged outcome is the sustained increase of this splicing enhancement over time, concurrently with the absence of cell toxicity or compromised activity, thereby affording new avenues for dosage optimization and the control of ASO stability. Thus, an experimental blueprint was devised to scrutinize the potential of XNA-DNA mixmer ASOs (MERLiNA and GENLiNA), which were tailor-designed to classic ISS-N1 targeted RNA-ASO sequences **(Figure 1)**.

**Figure 1:**
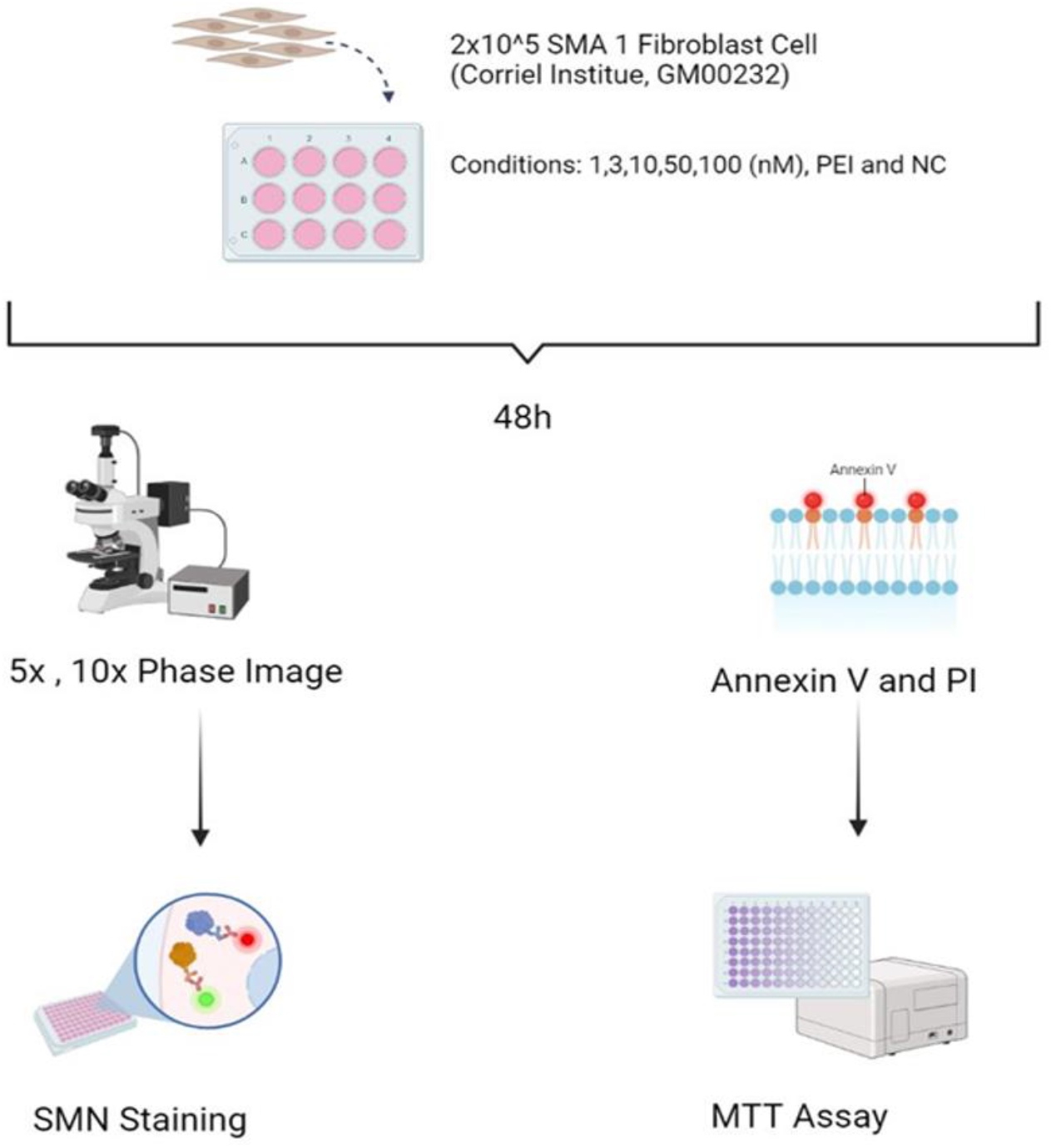
Experimental set-up for post-transfection analysis of XNA-DNA ASO mixmers on SMA type 1 primer fibroblast.

Preceding the analysis of XNA-DNA ASO mixmers, an inquiry into the optimal dose and SMN expression–cell viability balance of ASOs was pursued. To this end, a range of doses (1-200 nM) of PEI-coated RNA-ASO positive control were transfected into SMA type 1 fibroblast cells, with the consequent impact on SMN expression and viability being juxtaposed for comparative assessment. Notably, a discernible elevation in SMN expression emerged at 1nM from the 24-hour mark, with the most remarkable augmentation realized at 200 nM **(Figure 2a)**. Conversely, while SMN expression between 1-100 nM exhibited observable trends, it failed to manifest a significant deviation when compared to untreated and PEI-transfected controls **(Figure 2a)**. Remarkably, though 200 nM transfections incited an increase in SMN expression, cell death rates were markedly mitigated at 100 and 200 nM doses vis-à-vis the control groups **(Figure 2b)**. These findings underscore the importance of commencing SMN expression analysis 48 hours post-transfection in future studies, while emphasizing the comparison of concentrations below 100nM.

**Figure 2:**
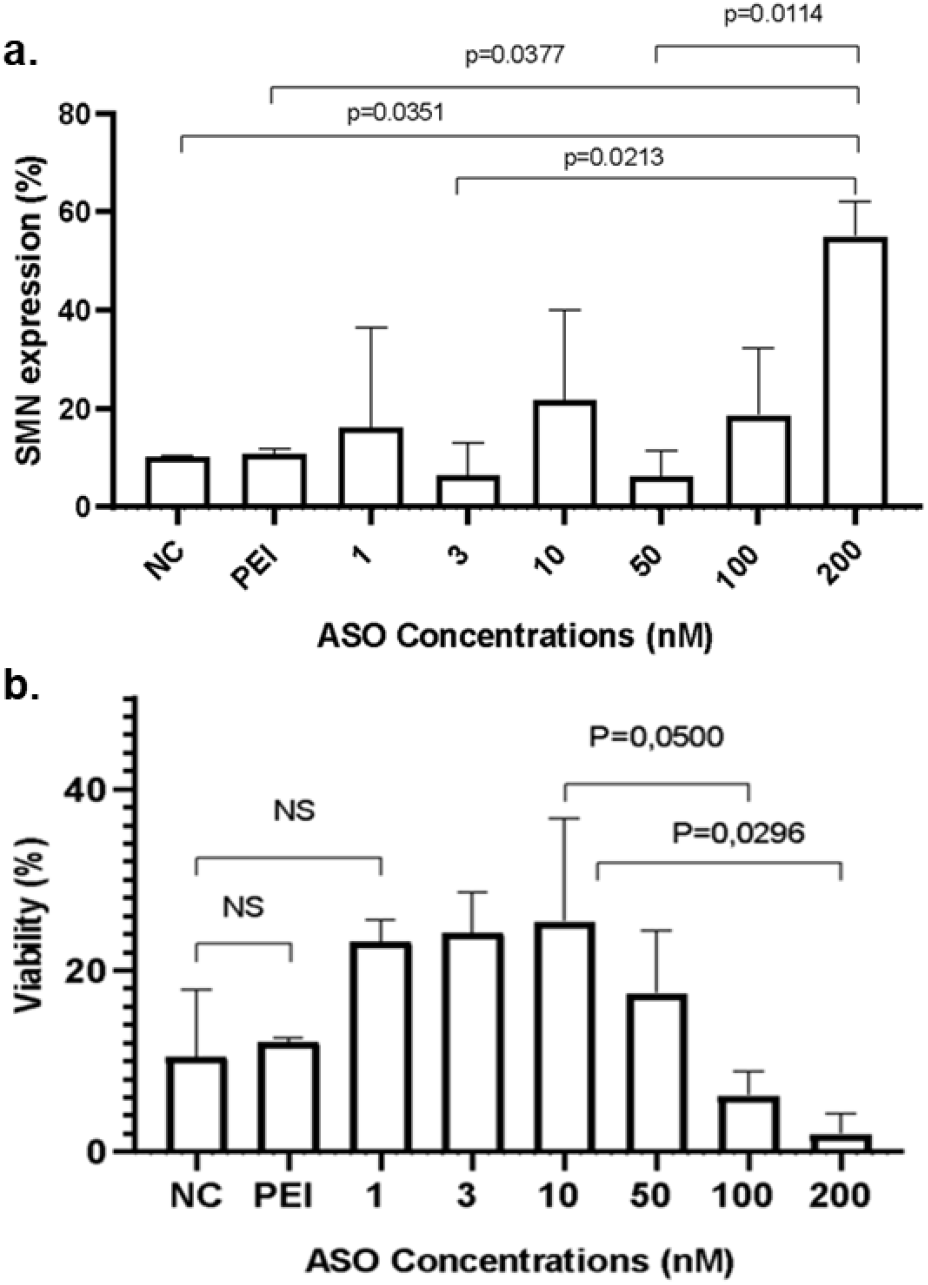
Upregulation of SMN protein expression determined as Viability. **a**. 24th hour analysis of SMN expression by flow cytometry of negative control SMA fibroblast cells, PEI transfected controls, and positive control ASO-treated SMN antibody staining at concentrations of 200nM, 100nM, 50nM, 10nM, 3nM and 1nM. **b**. Bar graph showing, 24th hour analysis of cell viability by flow cytometry of negative control SMA fibroblast cells, PEI transfected controls, and positive control ASO-treated SMN antibody staining at concentrations of 200nM, 100nM, 50nM, 10nM, 3nM and 1nM. (P < 0.05 is significant NS= non-Significant.)

Against this backdrop, positive control RNA-ASO, alongside GENLiNA and MERLiNA mixmer sequences, were interfaced with SMA type 1 fibroblast cells via PEI-mediated transfection. Post 48 hours, the first tier encompassing cell viability and SMN expression was interrogated through flow cytometric assessment subsequent to intracellular staining **(Figure 3a and 3b)**. Further delving into cellular dynamics, Annexin V and PI analysis coupled with flow cytometry was deployed to ascertain the induction of apoptosis pathways in cells, elucidating both early and late apoptosis phases **(Figure 3c)**. As an ancillary facet, cellular morphology was observed through inverted microscopy **(Figure 3d)**. Preliminary insights divulged that 1nM MERLiNA and GENLiNA conferred analogous or elevated SMN expression in comparison to the RNA-ASO positive control group **(Figure 3)**. Notably, the relatively low proportion of cells initiating the apoptosis pathway in contrast to the positive control and PEI-transfected control groups suggested the favorable influence of GENLiNA and MERLiNA on cell viability. Importantly, these findings, showcased in **Figure 3**, were juxtaposed against untreated and PEI-transfected control groups, laying the foundation for subsequent quantification in the flow analysis.

**Figure 3:**
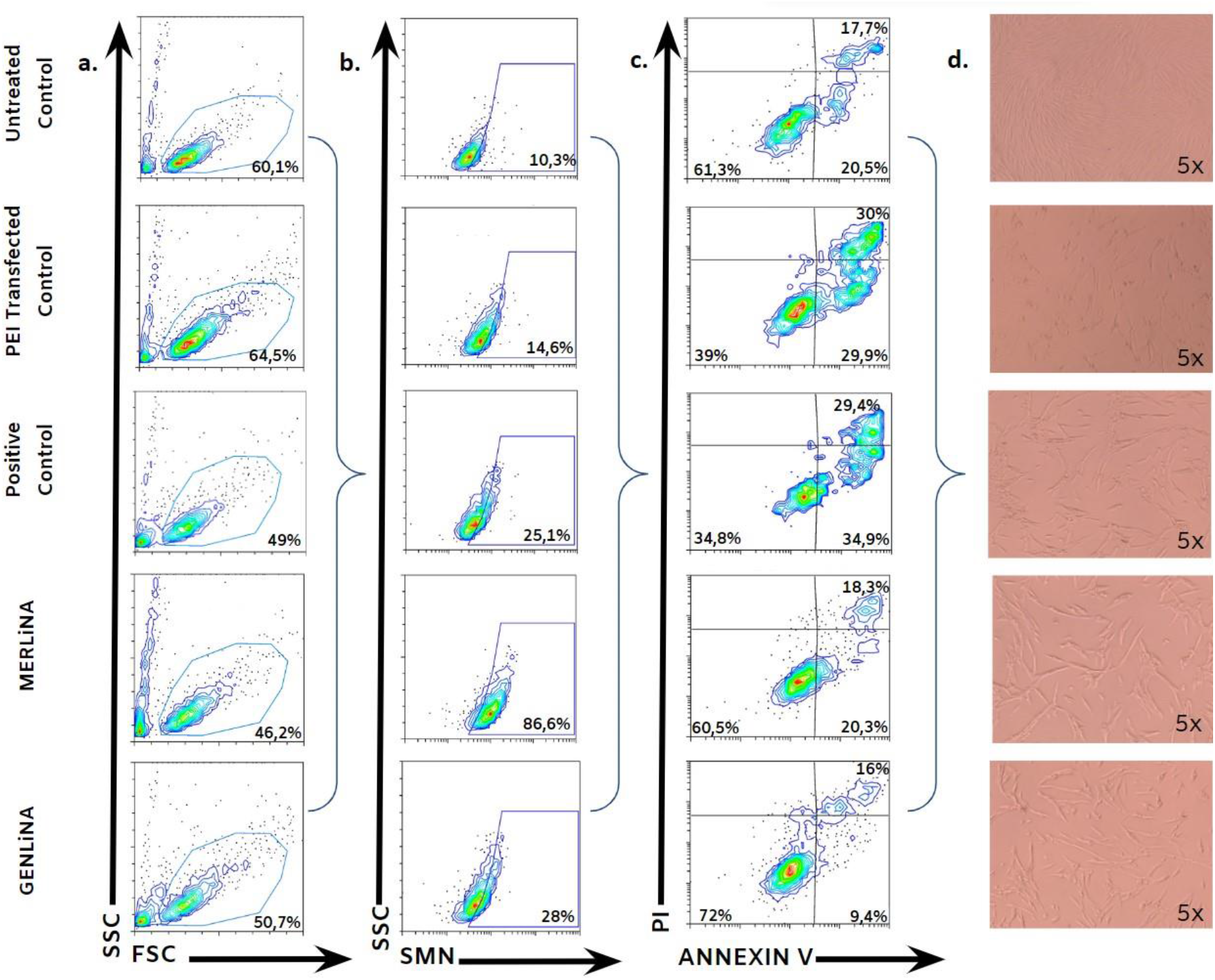
Flow Cytometry Analysis of ASO sequences transfected with PEI. **a**. 48th hour analysis of cell viability of SMA Fibroblast cells in flow cytometry treated with untreated control, PEI transfected control, positive control (1nM), MERLiNA (1nM) and GENLiNA (1nM) **b**. 48th hour analysis of SMN expression in flow cytometry of SMN antibody staining of SMA fibroblast cells treated with untreated control, PEI transfected control, positive control (1nM), MERLiNA (1nM) and GENLiNA (1nM). **c**. 48th hour flow cytometry analysis of Annexin V-PI antibody staining of SMA fibroblast cells treated with untreated control, PEI transfected control, positive control (1nM), MERLiNA (1nM) and GENLiNA (1nM). **d**. 48th hour 5x phase images of SMA fibroblast cells under fluorescent microscope, treated with untreated control, PEI transfected control, positive control (1nM), MERLiNA (1nM) and GENLiNA (1nM).

Following meticulous optimizations spanning both dose and temporal aspects, RNA-ASO positive control, coupled with GENLiNA and MERLiNA XNA-DNA ASO mixmer groups, were translocated onto SMA type 1 fibroblast cells in a dose gradient of 1-100 nM over a 48-hour duration. The outcomes reflected a noteworthy escalation in SMN expression across all dosage levels vis-à-vis control groups (NC and PEI), signifying that these augmentations didn’t compromise cell viability **(Figure 4a and 4b)**. Particularly, the viability of GENLiNA and MERLiNA, especially at doses of 50 and 100 nM, outshone the positive control group **(Figure 4a)**. Intriguingly, the 1nM MERLiNA dose rendered a significantly greater SMN expression than the GENLiNA group and even outperformed the positive control **(Figure 4b)**. This panorama underscores the ability of MERLiNA and GENLiNA XNA-DNA ASO mixmers to match, or even surpass, the SMN expression levels induced by the classical RNA-ASO positive control group, while concurrently showcasing a superior cell viability profile.

**Figure 4:**
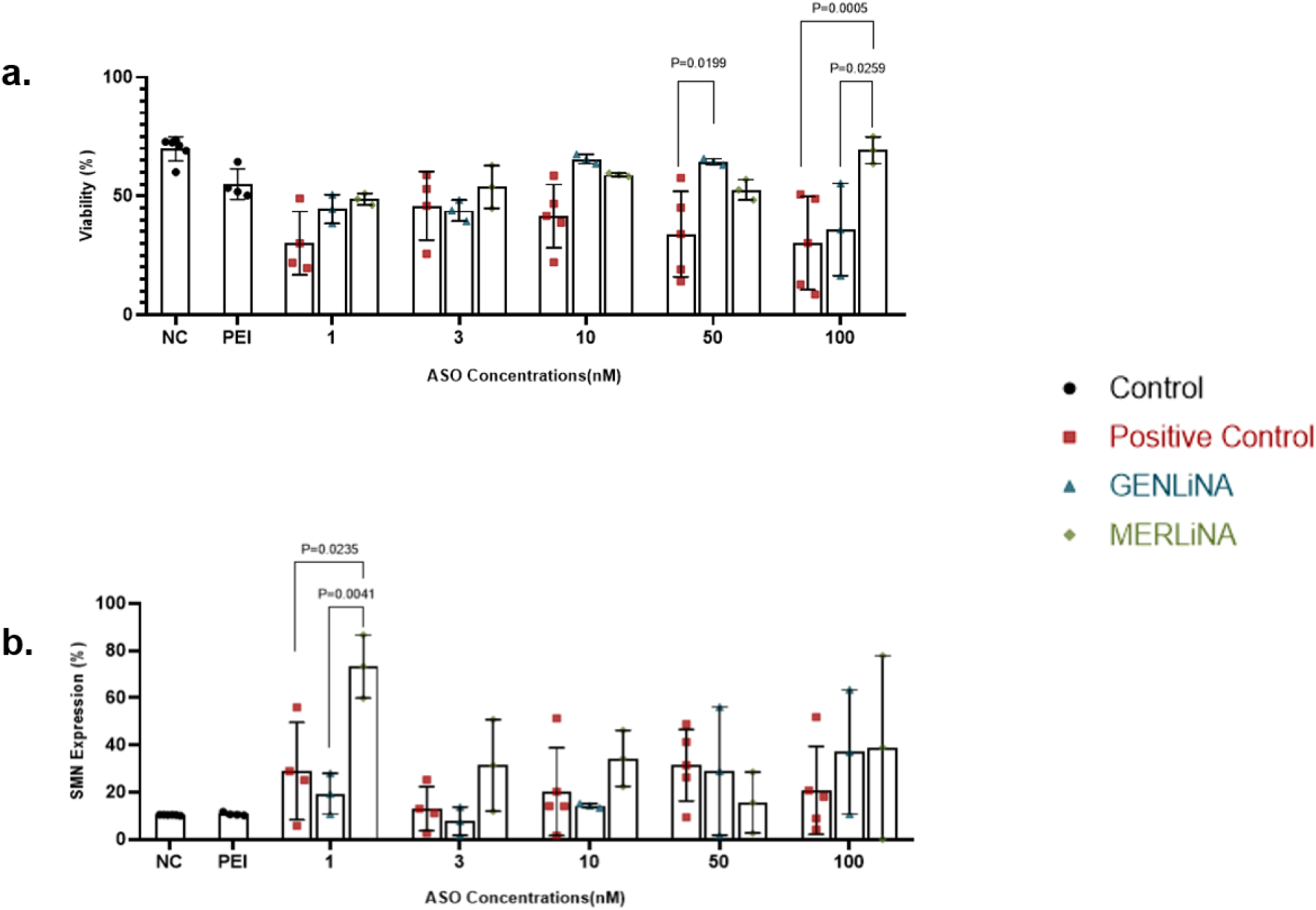
SMN protein expression vs Viability upon post-transfection of ASOs. **a**. Bar graph showing, 48th hour analysis of cell viability of SMA Fibroblast cells in Untreated Control, PEI Control, Positive Control, MERLiNA and GENLiNA. **b**. Bar graph showing, 48th hour analysis of SMN expression of SMA Fibroblast cells in Untreated Control, PEI Control, Positive Control, MERLiNA and GENLiNA. (Black is control, red is Positive Control, blue is GENLiNA, green is MERLiNA). (P < 0.05 is significant).

The subsequent facet of the investigation, centered on a detailed analysis of viability within these enhanced SMN expression dose groups, hinged upon the measurement of early and late apoptosis rates via Annexin V and PI staining, 48 hours post-ASO application. These deliberations revealed that MERLiNA administration correlated with heightened cell viability across numerous dose groups compared to the positive control and GENLiNA **(Figure 5a)**. This advantageous effect was concurrently mirrored in the reduced incidence of early and late phase apoptosis in the MERLiNA group as opposed to these comparators **(Figure 5b and 5c)**. The collective implications of these findings point toward MERLiNA’s potential to confer improved cell viability in vitro relative to the classical RNA-ASO positive control group.

**Figure 5:**
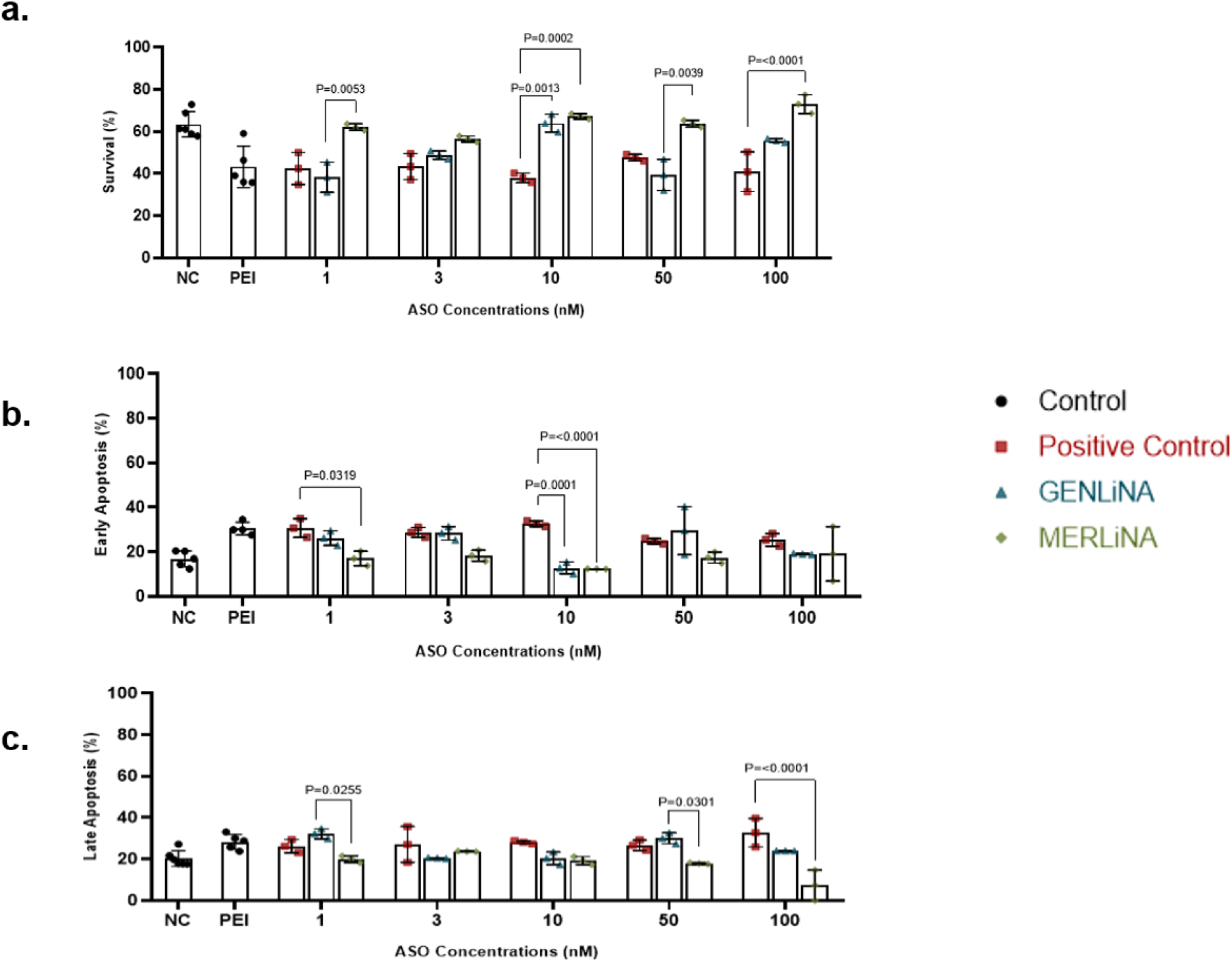
Apoptotic effect at 48th hour after transfection with ASO sequence in primary SMA fibroblast cell line. **a**. Bar graph showing the proportion of Annexin-V negative PI negative viable cells after annexin-V/PI staining. **b**. Bar graph showing the proportion of Annexin-V positive PI negative cells in the early apoptosis stage. **c**. Bar graph showing the proportion of Annexin-V positive PI positive cells in the late apoptosis stage. (Black is control, red is Positive Control, blue is GENLiNA, green is MERLiNA) (P < 0.05 is significant)

The investigation further delved into assessing how mitochondrial activity underwent transformation within populations of SMA type 1 fibroblast cells exposed to ASO groups, an analysis facilitated through MTT assessment, which gauges mitochondrial reductase activity. Notably, no significant downturn in mitochondrial activity was discerned in contrast to the NC and PEI control groups, underscoring that the GENLiNA and MERLiNA XNA-DNA ASO mixmers preserved cell viability relative to the NC, PEI, and RNA-ASO positive control groups **(Figure 6)**.

**Figure 6:**
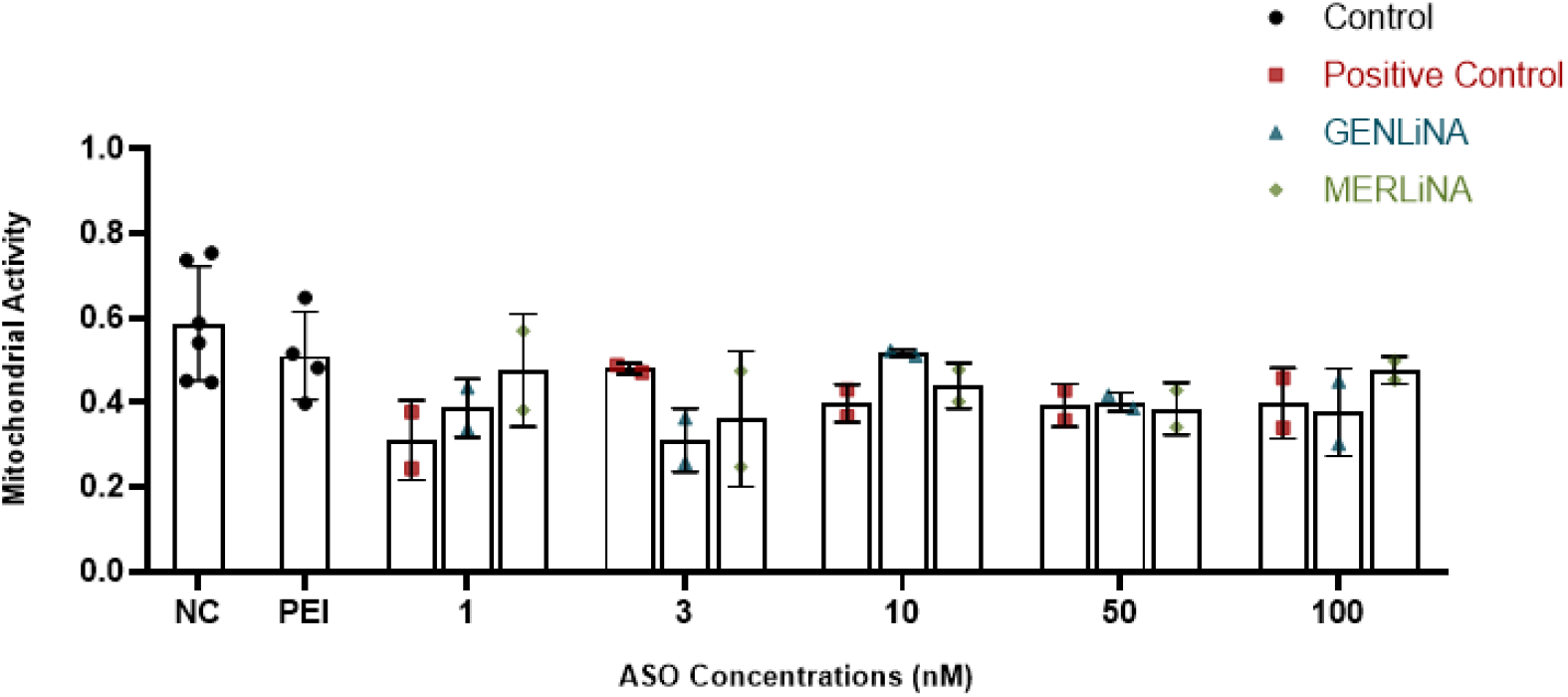
Mitochondrial activity (MTT) analysis. (Black is control, red is Positive Control, blue is GENLiNA, green is MERLiNA).

## Discussion

The outcomes of this study underscore the potential of XNA-DNA-ASO mixmers as a promising avenue for enhancing functional SMN protein expression in SMA type 1 fibroblast cells. The splicing modulation properties of ASOs have been previously demonstrated, particularly in the context of SMA, as evidenced by studies targeting the ISS-N1 region (Singh et al., 2013; Zhang et al., 2010). Our findings contribute to this body of knowledge by exploring the utilization of XNA-DNA-ASO mixmers, which offer distinct advantages due to their chemical composition and higher endonuclease resistance, as compared to classical RNA-ASOs. XNA polymers, with their modified sugar backbone, possess an enhanced binding affinity for RNA sequences and increased resistance to nuclease degradation (Wang et al., 2022; Tateishi-Karimata, 2020). This enhanced stability potentially contributes to the sustained effects observed in our study, as evidenced by the prolonged increase in SMN expression over 48 hours without causing significant cell toxicity.

The current treatment landscape for SMA relies heavily on costly gene therapy interventions, such as Zolgensma and Spinraza, which aim to provide short-term benefits but lack long-term stability (Hoy, 2019; Mendell et al., 2017; Sivanesan, 2013). In contrast, the XNA-DNA-ASO mixmers investigated here offer a potential solution for sustained and stable enhancement of SMN protein expression. The effectiveness of ASO-based approaches, such as XNA-DNA-ASOs, is underscored by their ability to modulate splicing mechanisms, particularly those involving the critical exon 7 of the SMN2 gene. Our study builds upon previous research by demonstrating that XNA-DNA-ASOs effectively target the ISS-N1 region in intron 7, leading to increased exon 7 inclusion and subsequent SMN protein production.

Moreover, our investigation elucidated the critical balance between SMN expression enhancement and cell viability. The comparative analysis of different ASO doses revealed that both MERLiNA and GENLiNA XNA-DNA-ASO mixmers can achieve SMN expression levels comparable to, or exceeding, those induced by classical RNA-ASO treatment. Remarkably, these increased SMN expression levels were accompanied by improved cell viability, as evidenced by lower apoptotic rates and preserved mitochondrial activity. This dual effect suggests that XNA-DNA-ASOs not only enhance the production of functional SMN protein but also provide a more favorable cellular environment for viability.

Nevertheless, while our study highlights the promising potential of XNA-DNA-ASOs, further research is required to unravel the underlying molecular mechanisms governing their effects on splicing and stability. Moreover, in vivo studies and clinical trials will be imperative to validate the translational potential of XNA-DNA-ASOs as a safe and effective therapeutic strategy for SMA.

In conclusion, the results of this study contribute to the ongoing quest for improved therapeutic interventions for SMA. By harnessing the advantageous properties of XNA-DNA-ASOs, we present a compelling case for the sustained enhancement of SMN protein expression and cellular viability. These findings open up new avenues for the development of next-generation genetic therapies that offer long-term stability and improved outcomes for individuals affected by SMA.

## Notes

**Conflict of interest:** O.K., H.A.B., C.C.I., S.P.O., G.Y., C.T. are inventors of patent applications (pending) including “Methods For The Treatment Of Spinal Muscular Atrophy” (2021/018884) at Turkish Patent and Trademark Office. No other author has a competing interest except for these authors.

**Funding statement:** All funding in the study was supported by Üsküdar University Scientific Research Studies with the grant number, ÜÜBAP-YP-2021-014.

### Competing Interest Statement

O.K., H.A.B., C.C.I., S.P.O., G.Y., C.T. are inventors of patent applications (pending) including Methods For The Treatment Of Spinal Muscular Atrophy (2021/018884) at Turkish Patent and Trademark Office. No other author has a competing interest except for these authors.

